# Subchronic exposure to ambient PM_2.5_ impairs novelty recognition and spatial memory

**DOI:** 10.1101/2023.09.07.556582

**Authors:** Sun-Hong Kim, Darpan Das, Fenna C. M. Sillé, Gurumurthy Ramachandran, Shyam Biswal

**Author notes:** Correspondence: Shyam Biswal Gurumurthy Ramachandran Fenna Sillé.

## Abstract

Air pollution remains a great challenge for public health, with the detrimental effects of air pollution on cardiovascular, rhinosinusitis, and pulmonary health increasingly well understood. Recent epidemiological associations point to the adverse effects of air pollution on cognitive decline and neurodegenerative diseases. Mouse models of subchronic exposure to PM_2.5_ (ambient air particulate matter < 2.5 µm) provide an opportunity to demonstrate the causality of target diseases. Here, we subchronically exposed mice to concentrated ambient PM_2.5_ for 7 weeks (5 days/week; 8h/day) and assessed its effect on behavior using standard tests measuring cognition or anxiety-like behaviors. Average daily PM_2.5_ concentration was 200 µg/m^3^ in the PM_2.5_ group and 10 µg/m^3^ in the filtered air group. The novel object recognition (NOR) test was used to assess the effect of PM_2.5_ exposure on recognition memory. The increase in exploration time for a novel object versus a familiarized object was lower for PM_2.5_-exposed mice (42% increase) compared to the filtered air (FA) control group (110% increase). In addition, the calculated discrimination index for novel object recognition was significantly higher in FA mice (67 %) compared to PM_2.5_ exposed mice (57.3%). The object location test (OLT) was used to examine the effect of PM_2.5_ exposure on spatial memory. In contrast to the FA-exposed control mice, the PM_2.5_ exposed mice exhibited no significant increase in their exploration time between novel location versus familiarized location indicating their deficit in spatial memory. Furthermore, the discrimination index for novel location was significantly higher in FA mice (62.6%) compared to PM_2.5_ exposed mice (51%). Overall, our results demonstrate that subchronic exposure to higher levels of PM_2.5_ in mice causes impairment of novelty recognition and spatial memory.

## Introduction

Air pollution, a complex mixture of particulate matter, gases, metals and organic compounds, remains a major environmental threat to public health. We and others have demonstrated that ambient and indoor fine particulate matter smaller than 2.5 μm in diameter (PM_2.5_) plays an critical role in causality of cardiovascular, pulmonary, rhinosinusitis and metabolic diseases [1]. Interestingly, there is accumulating evidence that particulate matter can enter the brain [2–4]. In particular, ultra-fine particles which are smaller than 100 nm in diameter can enter olfactory bulb brain area through the translocation from the direct deposits at olfactory mucosa of the nasopharyngeal region of the upper respiratory tract (URT) [2–4]. Furthermore, these fine particles also can be reached to the central nervous system (CNS) through the lower respiratory tract (LRT) by crossing the impaired blood-brain-barrier [2]. Unlike our understanding of adverse effects of PM_2.5_ on respiratory and cardiovascular systems [5–7], the causal effect on neurodegeneration and brain diseases is unclear. Epidemiological studies have shown that chronic exposure to air pollution is associated with cognitive decline and increased neurodegeneration [8–11]. Indeed, PM_2.5_ particles were found deposited in various brain regions including cortical areas in frontal lobe, olfactory bulb and cerebellum [2–4]. Long-term exposures of concentrated ambient PM_2.5_ caused oxidative stress and neuroinflammation in CNS leading to various noxious outcomes such as neurodevelopmental impairment, neuronal loss and abnormal activation of sympathetic neurons leading to high blood pressure [12–15]. It is challenging to demonstrate causality of adverse effects of air pollution on the brain in human studies. Thus, studies with exposure of animal models to real-world PM_2.5_ can improve our understanding of causality and mechanisms. Here, we employed subchronic exposure of concentrated ambient PM_2.5_ to generate an appropriate mouse model and hypothesized that these exposures impair certain cognitive function, especially novelty recognition and spatial memory and affective behavior, especially anxiety-related behavior in mice after a relatively short period (7-weeks) of PM_2.5_ subchronic exposure.

## Methods

### Animal Model

All procedures were approved by the Institutional Animal Care and Use Committee at Johns Hopkins University. Wild-type mice (C57BL/6J) were purchased from the Jackson Laboratory. All mice were housed in the animal housing facility at the Johns Hopkins University School of Public Health under a controlled environment (target temperature of 22.2°C; target humidity of 42%; 14.5h/9.5h light/dark cycle - 6:30 a.m. on/9:00 p.m. off) and given *ad libitum* access to normal food (LabDiet, 5V12 - Pico-Vac® Verified 75 IF EXT) and water throughout the study. All mice used in this study were male mice of same age (∼ 1 month-old at the beginning of their PM_2.5_ exposure).

### Exposure model

The PM_2.5_ exposures were provided to the animals in the chambers as follows: 5 days per week (Mon-Fri) and 9 am to 5 pm (8 h/day) daily for 7 weeks. These animals were acclimated in our housing facility for 1 week before the PM^2.5^ exposure and they were around 1 month-old at the beginning of PM_2.5_ exposure. We included 10 mice per each group [filtered air (FA) or PM_2.5_ (PM). Each group of mice (FA or PM) were regularly switched between the two separate exposure chambers (5 mice/chamber; 2 separate chambers for each FA or PM group of mice) throughout 7-week exposure period. There were no differences in locomotor activities or mobility in any of the mice assigned across groups prior to or during their exposure. Behavioral studies were performed at the end of the 7-week exposure period.

A Concentrated Ambient Particles System (CAPS) (Fig. 1) was developed to conduct subchronic animal inhalation studies on the health effects of air pollution extending over many months. The system uses the principle of the condensational growth of the ambient particles followed by virtual impaction to concentrate the aerosol [16]. Ambient aerosol was drawn at a flow rate of 1000 L/min through a cyclone inlet that removes most of the particles larger than 2.5 µm in aerodynamic diameter. The remaining fraction is representative of the urban ambient aerosol exposure in Baltimore, as shown in a previous detailed particle size distribution analysis of Baltimore aerosol, indicating a typical bi-modal number distribution representative of urban aerosol characteristics, with most of particles less or equal to 0.1μm in aerodynamic size [17]. A portion of the cyclone outflow (440 L/min) was passed over the warm bath of water (70°C). The saturated aerosol then entered the condenser (Model AP15R-40-A11B, PolyScience Inc., Niles, IL, USA), where it was rapidly cooled to about −7°C, resulting in supersaturation. The particles’ growth was induced by the condensation of water vapor onto the particle surface. The temperatures of water-bath vapor and the chiller were continuously monitored. Upon the exit of the condenser, the aerosol flow was split four ways, and the grown particles are concentrated using a virtual impactor operating at a major flow of 105 L/min and a minor flow of 5 L/min. Under ideal conditions, this should provide a concentration enrichment factor (CEF) of 20. The concentrated particles in the minor flow were passed through diffusion dryers to remove moisture and delivered to the exposure chambers. A portion of the major flow from the virtual impactors was passed through HEPA filters and was available for use for dilution of the exposure chamber concentration to the desired concentration. After dilution air was added, the system provided a CEF of 10-13 (Fig. 1B). See Figure 1A for a schematic of the system. For the sham control experiment, an identical system was used, except that a HEPA filter at the inlet to the mice chambers removed ambient particles and ensured the control mice receive clean air. The mice were placed in modular whole-body mouse exposure chambers that can accommodate 10 mice each and up to 4 cages were available for exposures at any given time (CH Technologies, Inc.). Aerosol exposures were monitored using a laser photometer (SidePak™ Personal Aerosol Monitor AM510, TSI, Inc., Shoreview, MN, USA). Figure 1B. shows the average daily PM concentrations in the exposure and sham (unexposed) chambers over a 24-day period.

**Figure 1.**
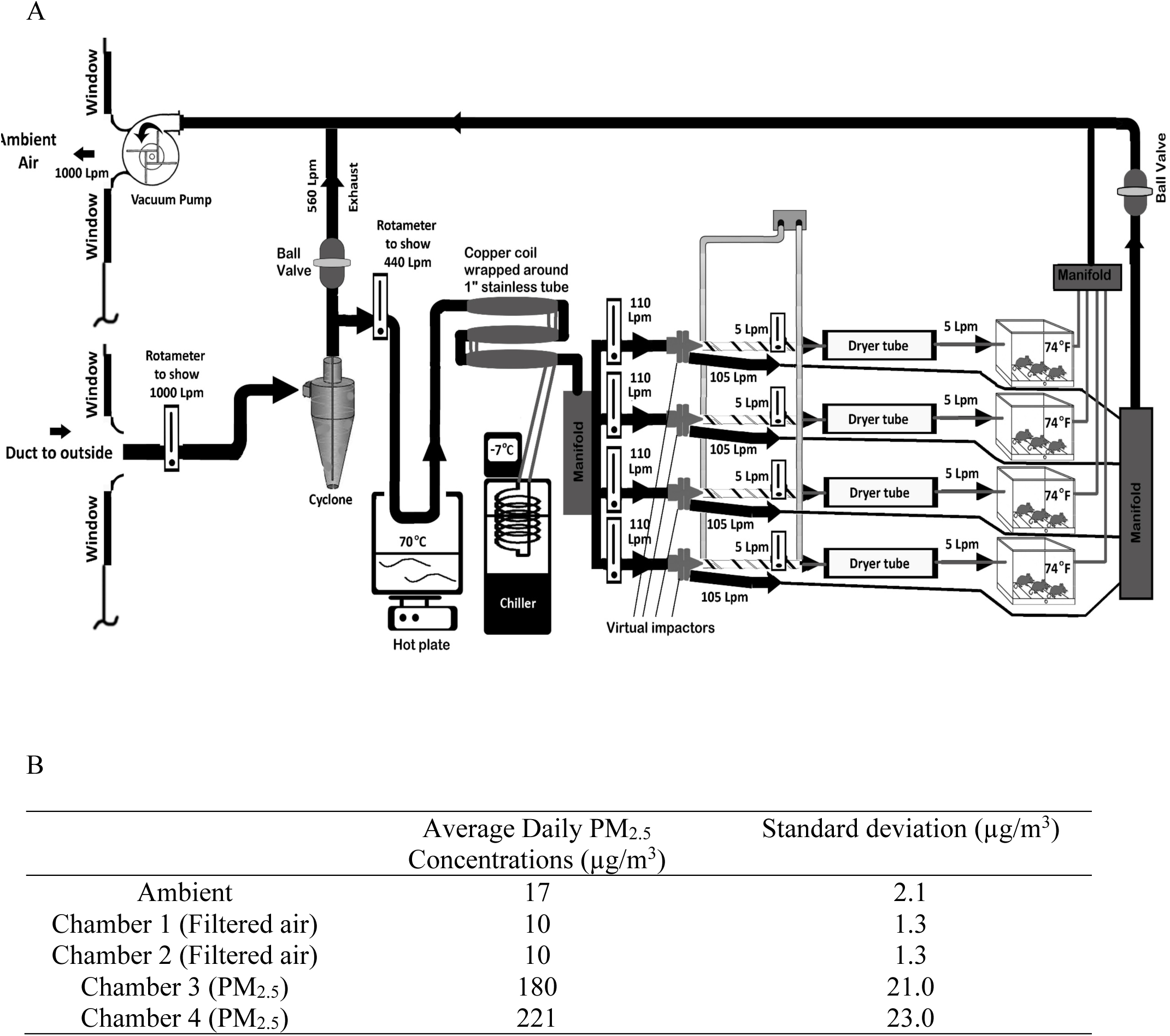
Exposure of concentrated ambient PM_2.5_ in mouse chambers. (A) Schematic of the Concentrated Ambient Particles System (CAPS). (B) Average daily PM_2.5_ concentrations (µg/m^3^) measured over 24 days in the Baltimore ambient air, and exposure (PM_2.5_) and sham (filtered air, FA) chambers.

### Behavioral tests

#### A novel object recognition task

(NOR) was used to assess recognition memory of a novel object as a representative form of cognitive function [18]. The NOR test has been widely used to examine novelty recognition memory using rodents. The apparatus consisted of a black-painted acrylic chamber (50 cm X 50 cm X 40 cm) with a video tracking system (CleverSys). The training session began by placing a mouse in the apparatus containing two identical objects and allowing it to explore for 10 min. After a 30 min time interval in its home cage, the test session began where the mouse was again introduced into the apparatus. But, in the test session, one of the two identical objects in the training session was replaced with a different object having a different shape and texture, and the mouse was allowed to explore for 5 min. The time spent exploring either the novel object or the familiarized object was counted to measure the cognitive performance of each mouse. Exploration was defined as directing the nose to the object and/or touching the object with the nose or forepaws.

#### An object location test

(OLT) was used to assess recognition memory of novel location [19]. The apparatus consisted of a black-painted acrylic chamber (50 cm X 50 cm X 40 cm) with a video tracking system (CleverSys). The training session began by placing a mouse in the apparatus containing two identical objects placed at two corners of the apparatus and allowing it to explore for 10 min. After a 30 min time interval in its home cage, the mouse was again introduced into the apparatus. In this test session, one object among two in the training session was placed in a different corner, and the mouse was allowed to explore for 5 min. The times spent exploring the novel location and the familiarized location were counted to measure cognitive performance of each mouse. Exploration was defined as directing the nose to the object and/or touching the object with the nose or forepaws.

#### Elevated plus maze

(EPM) was used to assess anxiety-like behaviors [20]. The maze consisted of two enclosed arms and two open arms with a video tracking system (CleverSys). A mouse was placed at the intersection of the open and closed arms, and the exploration time on the open arm was measured during a 5 min period. The maze was cleaned between tests with each animal.

We used MB10 (disinfectant) to clean the bottom of the chambers (NOR and OLT) and the maze (EPM) between tests. Briefly, before test began, the apparatus was sprayed with MB10 solution (Quip Laboratories, MBTAB6GRAM) and then wiped down completely. A subject animal was placed in the cleaned apparatus 30 sec after the final wipe. After one test, we removed visible urine and feces on the bottom with distilled water and sprayed with MB10. We waited 30 sec after wiping down of MB10 and then next subject animal was placed to the apparatus for new test.

### Statistical analysis

Statistical analysis was run using GraphPad Prism statistical software Version 9.5.1 (GraphPad Software, San Diego, CA). For the NOR and OLT data, we used parametric tests since the data displayed normal Gaussian distribution. Multiple paired *t*-tests were performed on paired data between familiar and novel (each mouse explored both familiar and novel) within each exposure group (FA and PM) and the results were corrected for multiple comparisons using the Holm-Šídák method resulting in adjusted p-values. Next, an unpaired two-sample parametric *t*-test comparing the FA and PM groups on these paired differences (novel -/- familiarized) within each group was performed to determine the effect of PM_2.5_ exposure on the delta (novel -/- familiarized) object exploration time compared to FA controls. In addition, the difference between FA and PM groups was determined with the mean discrimination index for novel object 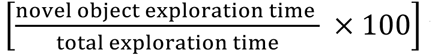 using a two-tailed unpaired parametric *t*-test.

For the Elevated plus maze (EPM) open arm (OA) time and OA entry, we used two-tailed unpaired parametric *t*-tests to determine the difference between the FA and PM exposure groups. In this whole study, we set *p* < 0.05 as the significant statistical value and described such significant ones as star (*)-labeled as shown in the figure legends (*p* < 0.05; **, *p* < 0.01; ***, *p* < 0.001). All data in this study is reported as Mean ± standard error of the mean (SEM).

## Results

To determine whether subchronic PM_2.5_ exposures affect cognitive function, we first tested two groups of mice that had been exposed to either filtered air or “real-world” highly concentrated PM_2.5_-containing air for 7 weeks using a Concentrated Ambient Particles System (CAPS) (Fig. 1). These mice were tested for their novelty recognition function using a novel object recognition (NOR) test where a mouse was presented with two identical objects during the first session, and then, after 30 min of interval time, one of the two objects was replaced by a new object during a second session (Fig. 2A). The amount of time taken to explore the new object provided an index of recognition memory. The group of mice exposed to filtered air (FA) spent a significantly longer time exploring the novel object compared to the already familiarized object (a 110 % increase, Fig. 2B and 2D, n = 10, *p adjusted* = 0.000300). In contrast, the mice exposed with subchronic, high concentration of ambient PM_2.5_ (PM_2.5_) exhibited a lower difference in exploration time between familiarized object versus novel object indicating their deficit in novelty recognition memory (a 42 % increase Fig. 2B and 2E, n = 10, *p* adjusted= 0.045202). A two-sample parametric *t*-test comparing the FA and PM groups on these paired differences (novel -/- familiarized) indicated that PM_2.5_ exposure significantly lowered the delta novel object exploration time compared to FA controls (*p* = 0.0431). The calculated discrimination index for novel object, 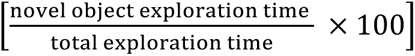 was significantly higher in FA mice (67 %) compared to PM_2.5_ exposed mice (57%) the latter which was close to random chance or 50% (Fig. 2C, *p* = 0.0190).

**Figure 2.**
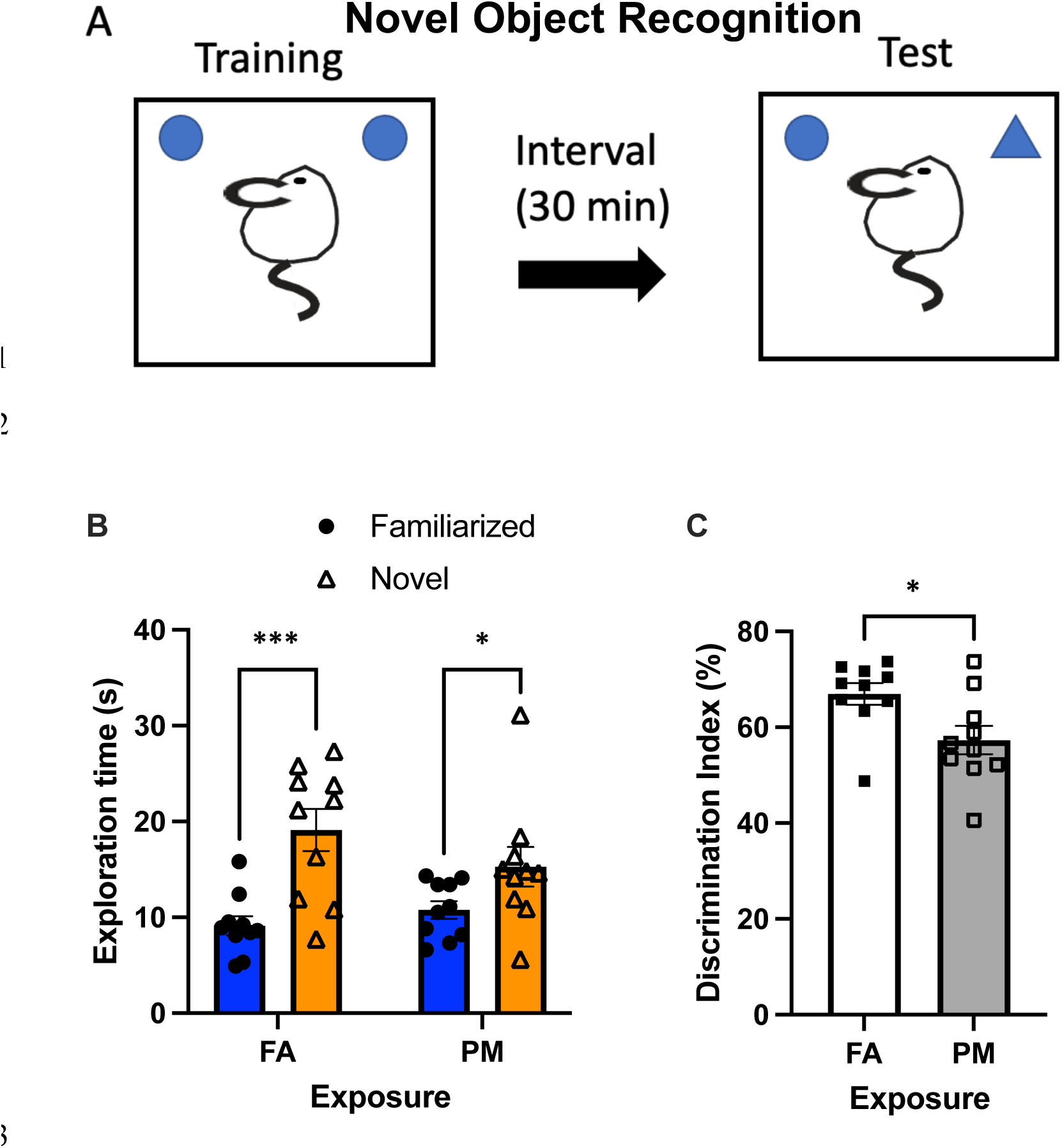
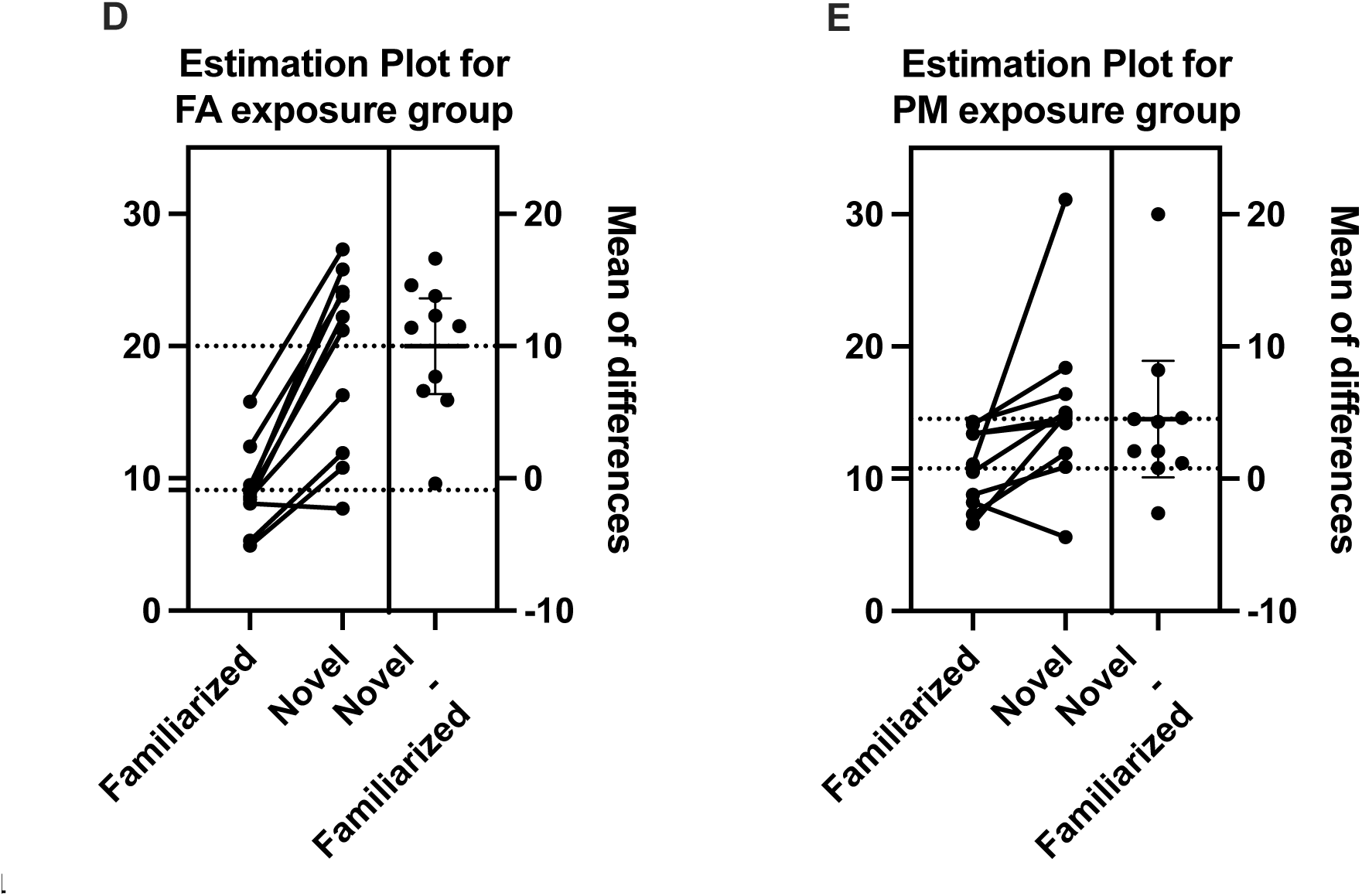
Adverse effect of PM_2.5_ exposure on recognition memory. (A) Scheme of novel object recognition task (NOR) with two objects. (B) Object exploration time. Impaired recognition memory in PM_2.5_ mice (n = 10, *p* adjusted= 0.045202) and normal novelty recognition performances in FA mice (n = 10, *p adjusted* = 0.000300). (C) Novel object discrimination index (DI). Significantly higher DI in FA mice (n = 10) compared to PM_2.5_ mice (n = 10), *p* = 0.0190. (D) Estimation plot indicating the mean of differences between the paired results from familiarized and novel object exploration time within the FA exposure group. (E) Estimation plot indicating the mean of differences between the paired results from familiarized and novel object exploration time within the PM exposure group. DI (%) = (Exploration time for novel object/Total exploration time) X 100. *, *p* < 0.05; ***, *p* < 0.001.

To confirm and support our finding about cognitive deficit caused by long-term ambient concentrated PM_2.5_ exposures, we also performed an object location test (OLT) which is another cognitive test using a different cognitive modality. While NOR examines novelty recognition memory, OLT examines spatial memory. In the experimental setup, a mouse was presented with two identical objects located at two corners of an arena during the first session, and after a 30 min time interval, one of the two objects was relocated to a different corner of an arena during a second session creating a novel space (Fig. 3A). The amount of time taken to explore the object relocated to a new location provided an index of spatial memory. FA mice spent a significantly longer time exploring an object at the novel location compared to an object at an already familiarized location (a 59.3 % increase, Fig. 3B and 3D, n = 10, *p* = 0.032698). However, the PM_2.5_ exposed mice exhibited no significant difference in their exploration time between familiarized location versus novel location indicating their deficit in spatial memory (a 3.7 % increase, Fig. 3B and 3E, n = 10, *p* = 0.723155). A two-sample parametric *t*-test comparing the FA and PM groups on these paired differences (novel -/- familiarized) indicated that PM_2.5_ exposure significantly lowered the delta object location exploration time compared to FA controls (*p* = 0.0382). Furthermore, the discrimination index for novel location was significantly higher in FA mice (62.6%) compared to PM_2.5_ exposed mice (51%) (*p* = 0.0314), the latter which was close to random chance or 50% (Fig. 3C). Together, our data provide supporting evidence that subchronic exposures of ambient concentrated PM_2.5_ can cause significant deficits in novelty recognition and spatial memory in mice compared to controls (FA).

**Figure 3.**
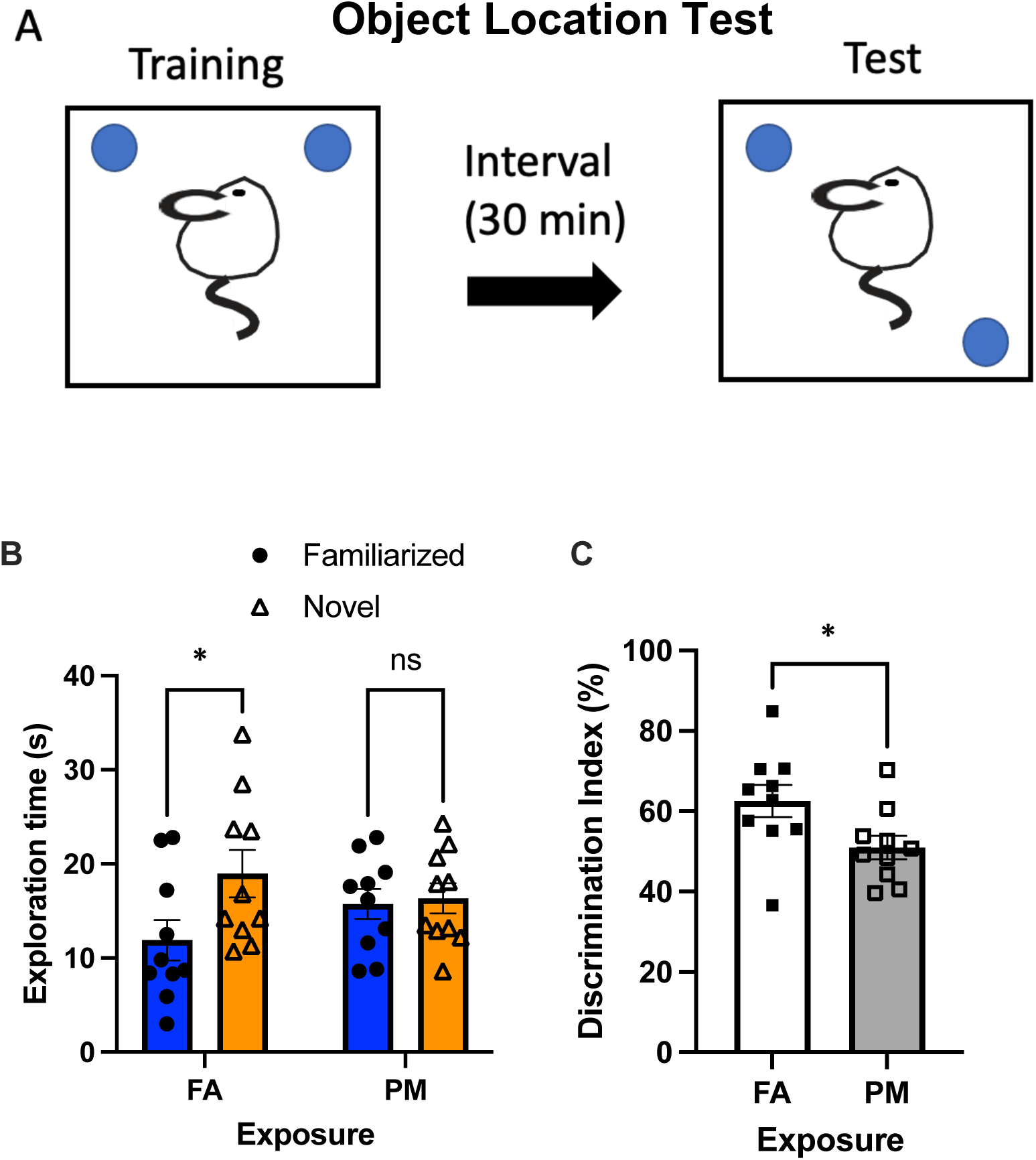
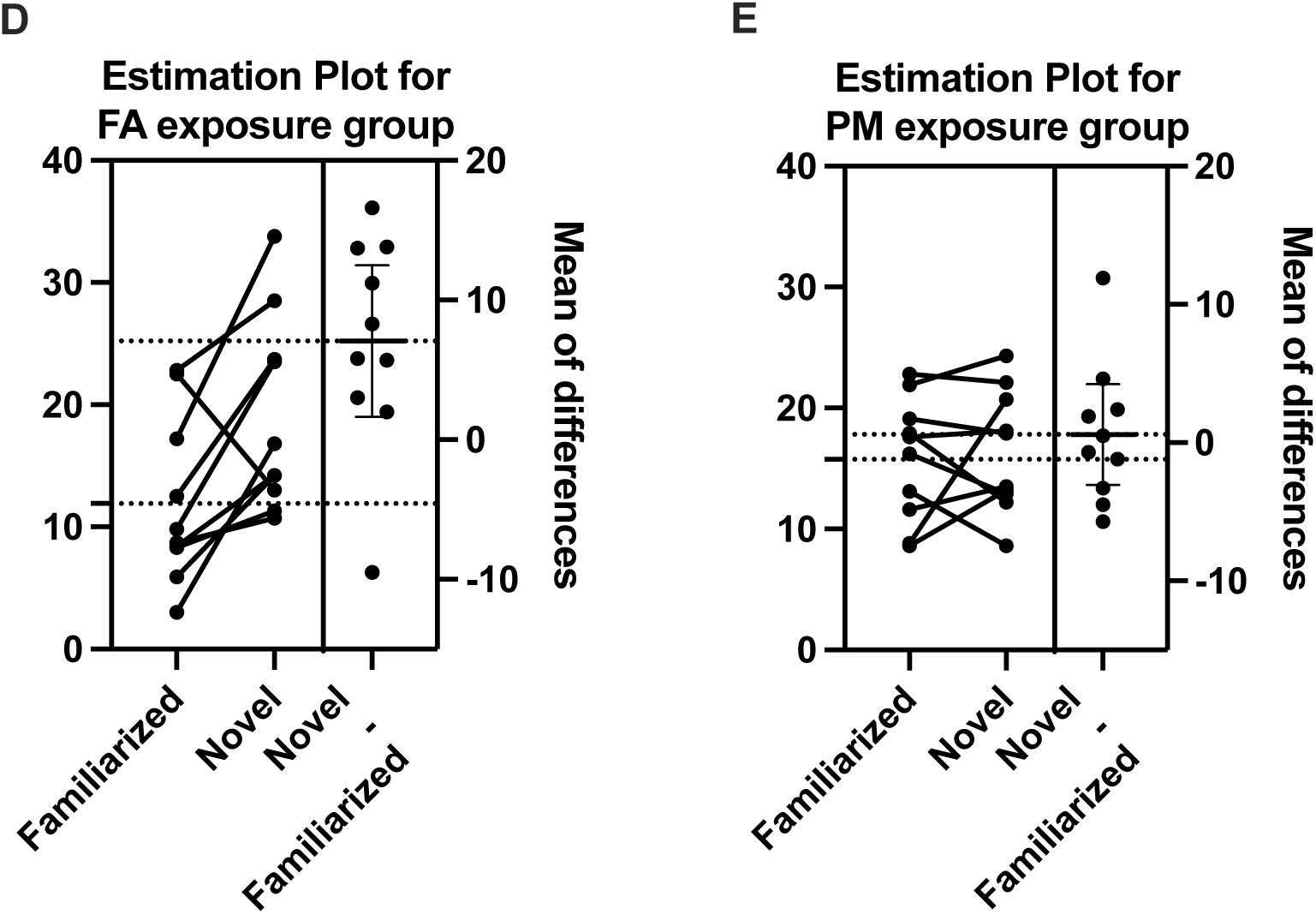
Adverse effect of PM_2.5_ exposure on spatial memory. (A) Scheme of object location test (OLT). (B) Object exploration time in PM_2.5_ mice (n = 10, *p* = 0.723155) and FA mice (n = 10, *p* = 0.032698). (C) Novel location discrimination index (DI). Significantly higher DI in FA mice (n = 10) compared to PM_2.5_ mice (n = 10), *p* = 0.0314. (D) Estimation plot indicating the mean of differences between the paired results from familiarized and novel location object exploration time within the FA exposure group. (E) Estimation plot indicating the mean of differences between the paired results from familiarized and novel location object exploration time within the PM exposure group. DI (%) = (Exploration time for novel object location exploration time/Total exploration time) X 100. *, *p* < 0.05.

To determine whether PM_2.5_ exposures can influence affective behaviors, we performed an elevated plus maze (EPM) test which is a standard test for measuring the level of anxiety-like behaviors. The amount of time taken to explore the open arms provided an index of anxiety-like behavior (Fig. 4A). PM_2.5_ exposed mice showed a decrease in the total exploration time on the open arm by 24.4 % compared to that of FA mice, however this data was not statistically significant (Fig. 4B, *p* = 0.085). The number of entries to the open arm was significantly fewer in PM_2.5_ exposed mice by 22.7% compared to FA mice (Fig. 4C; *p* = 0.0441).

**Figure 4.**
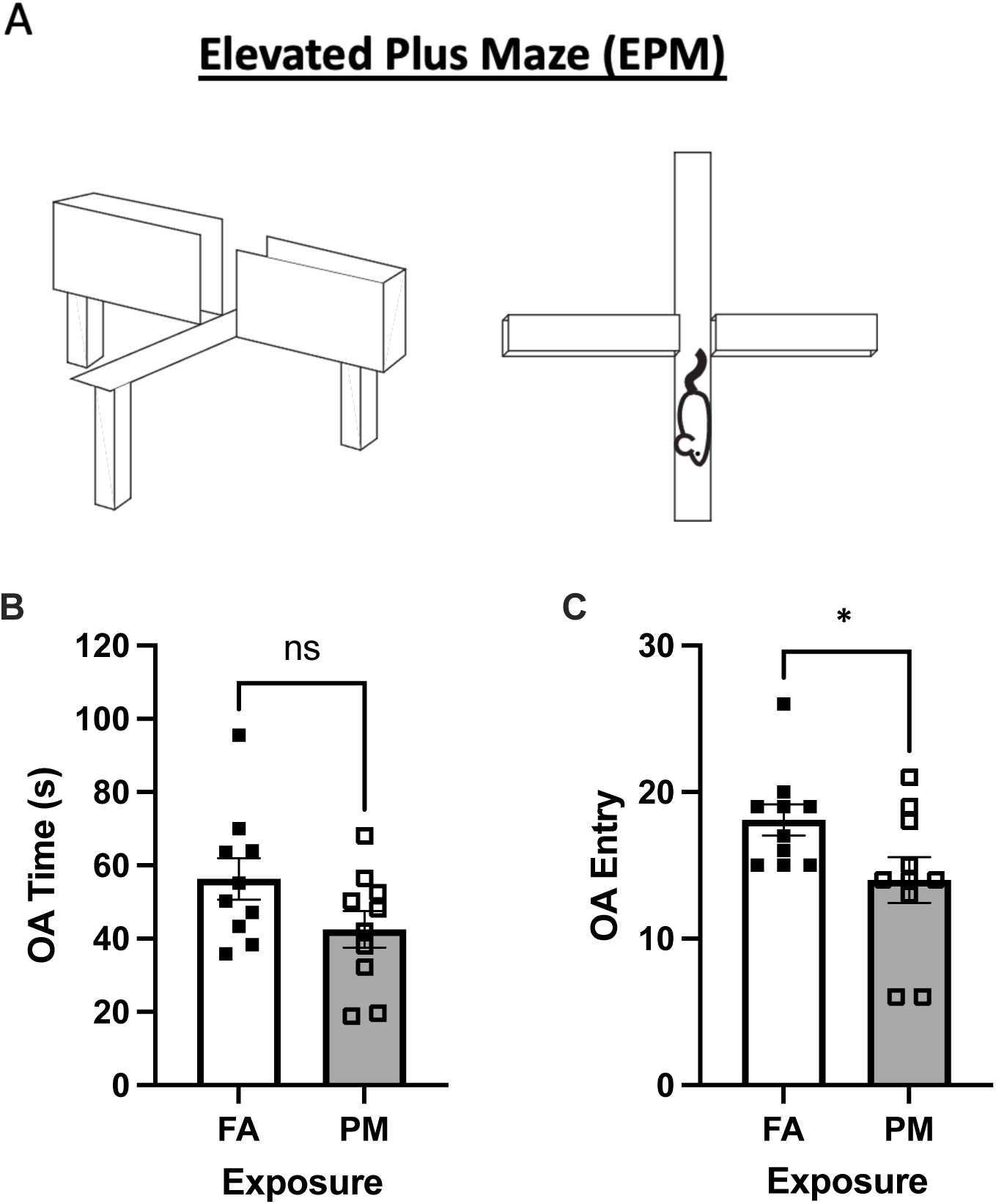
PM_2.5_-exposed mice did not show significant anxious behavior. (A) Elevated plus maze (EPM). (B) Open arm (OA) time. Decreased OA time in PM_2.5_ mice (n = 10; ns, *p* = 0.085) compared to FA mice (n = 10) indicating the anxious behavior caused by long-term PM_2.5_ exposure. (C) OA entry. PM_2.5_-exposed mice exhibited significantly fewer entry to open arms (n = 10; *p* = 0.044) compared to FA mice (n = 10). *, *p* < 0.05.

## Discussion

In this study, we showed the causality of long-term exposure (subchronic) of concentrated ambient PM_2.5_ in the impairment of cognitive functions such as recognition and spatial memory. While the contribution of specific brain regions and neuronal populations affected by PM_2.5_ is not yet clear, there is accumulating evidence that these fine particles can reach various brain regions including cortex and cerebellum through either URT or LRT route and result in neuroinflammation- and oxidative stress-mediated neuronal damages [2–4]. Although the mechanisms by which PM_2.5_ exposure might negatively affect brain function and PM_2.5_-caused behavioral deficits are largely unclear, chronic exposures of high concentration ambient PM_2.5_ have been linked to neuroinflammation, oxidative stress, neuronal loss in cortex and hippocampus and particle-induced disruption of the blood-brain barrier (BBB) in rodent and human brains [12,13,15,21–28]. Fagundes L.S. *et al*. also showed that the direct contact with fine particles increases oxidative stress in various brain regions, in particular hippocampus and cerebellum [29]. The hippocampus may be easily targeted by those fine particles entered CNS since hippocampus expresses abundant receptors for pro-inflammatory cytokines including interleukin (IL) 1β, IL6 and tumor necrosis factor α (TNFα) and thus can be particularly vulnerable to inflammation [13,15,30].

Long-term exposures to air pollution has been significantly associated with cognitive decline, increased neurodegeneration and major depressive disorder (MDD) in human epidemiological studies [8–11]. Similarly, controlled animal studies have revealed that long-term exposures to concentrated PM_2.5_ also cause cognitive decline and deficits in affective behaviors. However, mechanistic insights on the systemic effect of exposure to air pollution on the development of mental illnesses remain elusive. Since there are many different modalities in cognition, examination of cognitive functions determined by various concepts of cognition such as recognition, spatial, and social memory will help our better understanding of the adverse effects of subchronic PM_2.5_ exposures on cognition. In line with this idea, here we examined both novelty recognition and spatial memory with our concentrated ambient PM_2.5_-exposed animal model by performing the NORT test and OLT test, respectively. The cortex areas near hippocampus, perirhinal and entorhinal cortices, have been implicated as neural substrates for NORT, while the performance of OLT has been shown to be heavily rely on the hippocampal activities[31,32]. Since hippocampus and cortical areas near hippocampus has been known to be the target regions of fine airborne particles in the CNS, our behavioral data support the idea that prolonged exposure of PM_2.5_ results in cognitive decline likely due to hippocampal damage. Furthermore, here we report that a relatively much shorter period of exposure for only 7 weeks compared to other studies ranging 3 months to over 10 months also can cause significant cognitive deficit. This is particularly helpful information because we need to know the more accurate pathological kinetics caused by PM_2.5_ to better understand mechanisms underlying the pathophysiology of PM_2.5_. We also believe that the use of very young mice (∼ 1 month-old) for our PM_2.5_ exposure study likely contributed for the potentially accelerated occurrence of significant level of deficits in novelty recognition and spatial memory since postnatal to adolescent ages are very active period for brain structural development and thus are vulnerable to adverse factors that worsen cognitive function or emotional stability [33].

Chronic exposure of PM_2.5_ also has shown to increase the risk of disturbed emotional expression and related affective behaviors [34–36]. Previous studies have shown that long-term exposure of PM_2.5_ caused significant increase in anxiety-related behavior measured by EPM [37,38]. Although our EPM data (Fig. 4) showed some suggestive trend of anxious behavior, it did not reach statistical significance. EPM task has been shown to be heavily rely on the amygdala brain region, in particular basolateral amygdala (BLA) [39,40]. Our data may inform that the amygdala may be the brain region that is relatively less vulnerable to the negative effect of fine particles in CNS and likely need longer exposure to begin to show abnormality.

We suggest that the timing of apparent apperance of pathological signs caused by the prolonged exposure of PM_2.5_ can be variable depending on the PM_2.5_-targetted brain regions and their associated brain functions (e.g. cognition, anxiety, emotion, motivation, etc.) which may require different minimum duration of the exposure of PM_2.5_ needed to cause sufficient brain damages in that specific brain region. Thus, it would be interesting and informative to the better understanding of PM_2.5_-related pathophysiological kinetics if future studies examine and compare the physiological events (e.g. neuroinflammation or apoptosis) and molecular profiling (e.g. RNAseq or ATAC-seq) between the multiple groups of different duration of exposures of PM_2.5_ and groups of different ages at the beginning of exposure as well as between the different sexes. Detailed physical and chemical characterization of aerosol would give better insights on the PM_2.5_ associated health effects, and should be taken into account in future studies.

## Declarations

### Ethics

All procedures were approved by the Institutional Animal Care and Use Committee at Johns Hopkins University.

## Acknowledgments

We thank Kathryn A. Carson of the Johns Hopkins Biostatistics Center at the Bloomberg School of Public Health, Johns Hopkins University for consultation on statistical analysis of the data.

## Data availability statement

The data that support the findings of this study are available from the corresponding author, SB, upon reasonable request.

